# SuperPlotsOfData – a web app for the transparent display and quantitative comparison of continuous data from discrete conditions

**DOI:** 10.1101/2020.09.01.276881

**Authors:** Joachim Goedhart

**Affiliations:** Swammerdam Institute for Life Sciences, Section of Molecular Cytology, van Leeuwenhoek Centre for Advanced Microscopy, University of Amsterdam, P.O. Box 94215, NL-1090 GE Amsterdam, The Netherlands

## Abstract

Plots and charts are graphical tools that make data intelligible and digestible by humans. But the oversimplification of data by only plotting the statistical summaries conflicts with the transparent communication of results. Therefore, plotting of all data is generally encouraged and this can be achieved by using a dotplot for discrete conditions. Dotplots, however, often fail to communicate whether the data are from different technical or biological replicates. The superplot has been proposed to improve the communication of experimental design and results. To simplify the plotting of data from discrete conditions as a superplot, the SuperPlotsOfData web app was generated. The tool offers easy and open access to state-of-the-art data visualization. In addition, it incorporates recent innovations in data visualization and analysis, including raindcloud plots and estimation statistics. The free, open-source webtool can be accessed at: https://huygens.science.uva.nl/SuperPlotsOfData/

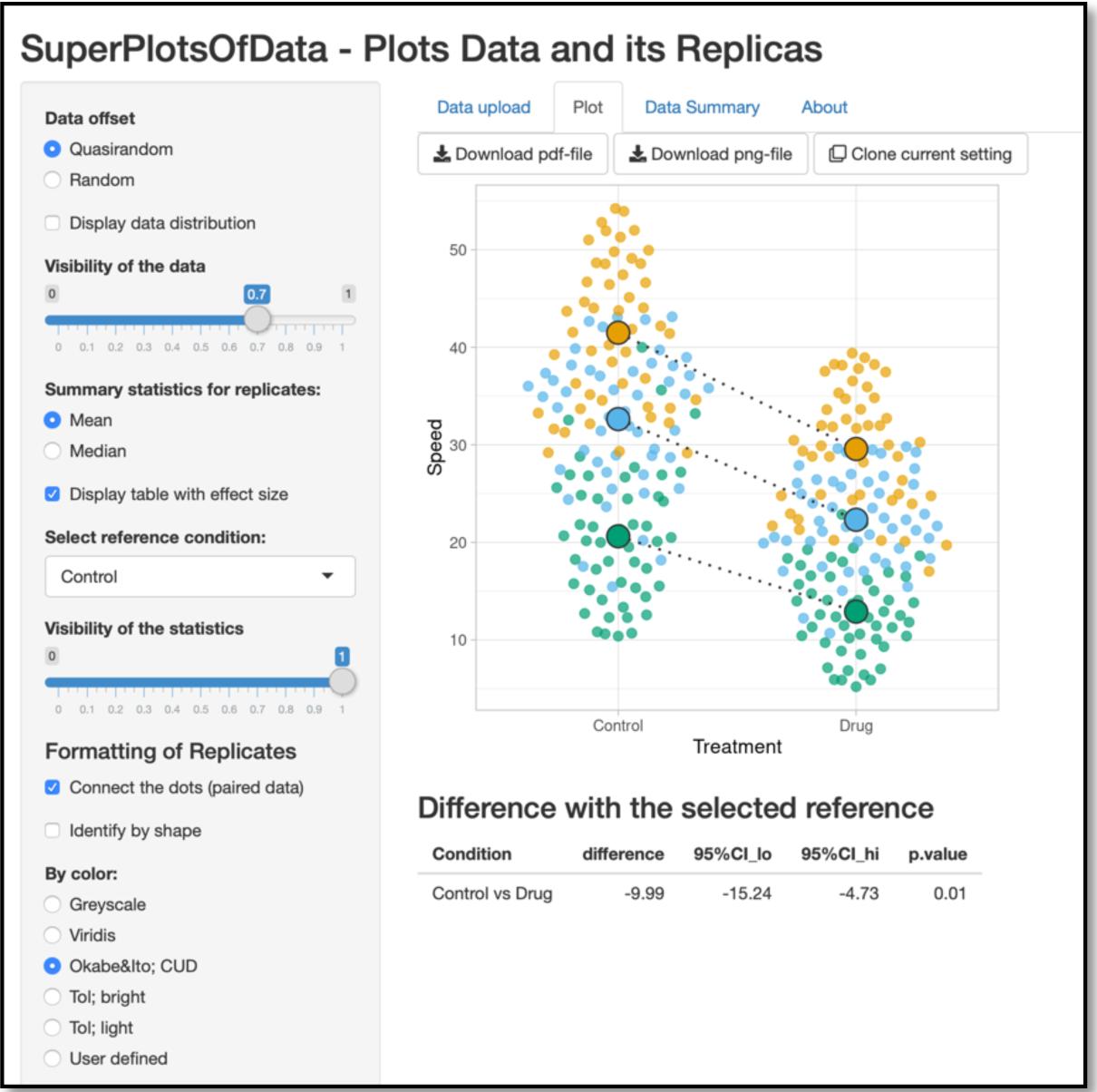

## Introduction

Graphs have a key role in the communication of experimental results in lab meetings, presentations and manuscripts. Over the years, efforts have been made to increase the transparency in reporting of results and this has led to the recommendation to plot all data, rather than just a summary (Weissgerber et al., 2019; Drummond and Vowler, 2011; Wilcox and Rousselet, 2018). For the display of continuous data from discrete conditions this means that instead of, or on top of, a bar that summarizes the data, the dot is the geometry of choice as it enables the display of all data. Dot plots can be generated by several popular commercial software packages, but there are also free, open source solutions (Weissgerber et al., 2017; Spitzer et al., 2014; Postma and Goedhart, 2019; Mauri et al., 2017).

We have previously created PlotsOfData to facilitate visualization of continuous data from different (discrete) conditions (Postma and Goedhart, 2019). Under the hood, PlotsOfData uses the statistical computing software R and several packages, including ggplot2 for state-of-the-art data visualization (Wickham, 2011). The software is operated by graphical user interface in a web browser. As a result, PlotsOfData is a universally accessible open source tool that delivers high quality plots, without the need for coding skills and with a minimal learning curve.

In addition to the transparent display of the data, information about the experimental design is necessary to interpret the results. For instance, it should be clear whether data are paired, how many technical and biological replicates are plotted and how ‘n’ is defined for different conditions (Lazic et al., 2018; Naegle et al., 2015). The ‘superplot’ has been proposed as a way to make this clear in a plot (Lord et al., 2020). A superplot identifies different replicates by color and/or shape and uses the average of each (biological) replicate as input for the comparison of conditions. An additional benefit of the explicit identification of biological replicates, instead of using the aggregated technical replicates, is that realistic p-values are obtained.

Other approaches that aim to improve data visualization and analysis have recently been proposed, i.e. raincloud plots (Allen et al., 2019) and the use of estimation statistics (Claridge-Chang and Assam, 2016; Cumming, 2014; Ho et al., 2019). The benefit of an open source tool (such as PlotsOfData) that is based on a powerful statistical computing language with excellent graphics is that these innovations are readily incorporated. In contrast, implementing these new ideas in commercial software requires multistep workarounds (Lord et al., 2020).

Here, SuperPlotsOfData is presented, which builds on the PlotsOfData web tool and uses the same philosophy of providing easy and open access to state-of-the-art data visualization. In addition, it enables the identification of replicates as a superplot and it incorporates recent innovations, including raindcloud plots and estimation statistics.

## Availability, code & issue reporting

The SuperPlotsOfData app is available at: https://huygens.science.uva.nl/SuperPlotsOfData The code was written using R (https://www.r-project.org) and Rstudio (https://www.rstudio.com). To run the app several freely available packages are required: shiny, ggplot2, magrittr, dplyr, readr, tidyr, ggbeeswarm, readxl, DT, broom and RCurl. This version of the manuscript is connected to version 1.0.1 of the web app (https://github.com/JoachimGoedhart/SuperPlotsOfData/releases/tag/v1.0.1), which is archived at zenodo, doi:10.5281/zenodo.4018039

Up-to-date code and new release will be made available on Github, together with information on running the app locally: https://github.com/JoachimGoedhart/SuperPlotsOfData/ The Github page of SuperPlotsOfData is the preferred way to communicate issues and request features (https://github.com/JoachimGoedhart/SuperPlotsOfData/issues). Alternatively, the users can contact the developers by email or <u>Twitter</u>. Contact information is found on the “About” page of the app.

## Data input and data format

The minimum input is a dataset with conditions in a column and the measured variables in another column. A third column that identifies the replicates is recommend to take full advantage of the application. This data format is generally known as tidy (Wickham, 2014). The data can be supplied as a CSV file, XLS(X) file, by copy-paste or through a URL (CSV files only). Two example datasets are available to demonstrate the data structure and for testing the app.

After data upload, the user selects the columns that hold the information on the Conditions, Measurements and (optional) Replicates. When no replicates are selected the measurements are grouped per condition.

## Data visualization

When the data is composed of different replicates, these are indicated for each condition with a different color and/or a different symbol. The mean or median of each of the replicates is indicated with a larger dot (figure 1). Lines are drawn between the means or medians from the same sample when the data is qualified by the user as ‘paired’. Pairing of the data will affect the statistics for the quantitative comparison of conditions, as will be explained below.

**Figure 1:**
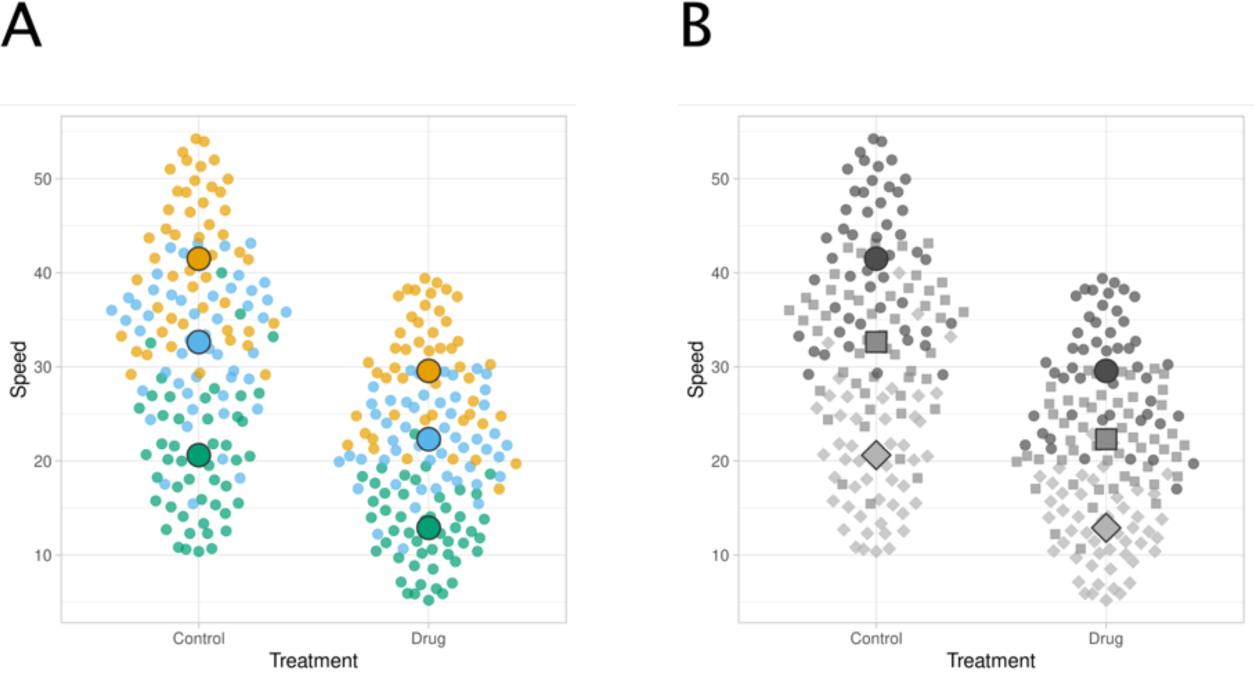
Output of the application based on example data. (A) By default, the (biological) replicates are identified by unique, colorblind friendly, colors and the mean of each replicate is indicated with a larger dot. (B) An alternative presentation of the same data that uses different symbols and grey values to identify replicates.

## Statistics

### Technical replicates

A median or mean can be selected as the summary statistic for each replicate and this is shown in the graphs as a larger dot. Several other statistics for individual replicates are available under the ‘Data Summary’ tab and include n, standard deviation and interquartile range (IQR). The table (figure 2) can be customized and by default lists the p-value from a Shapiro-Wilk test which can be used to evaluate whether the data of the replicates is normally distributed. A high p-value provides evidence for a normal distribution.

**Figure 2:**
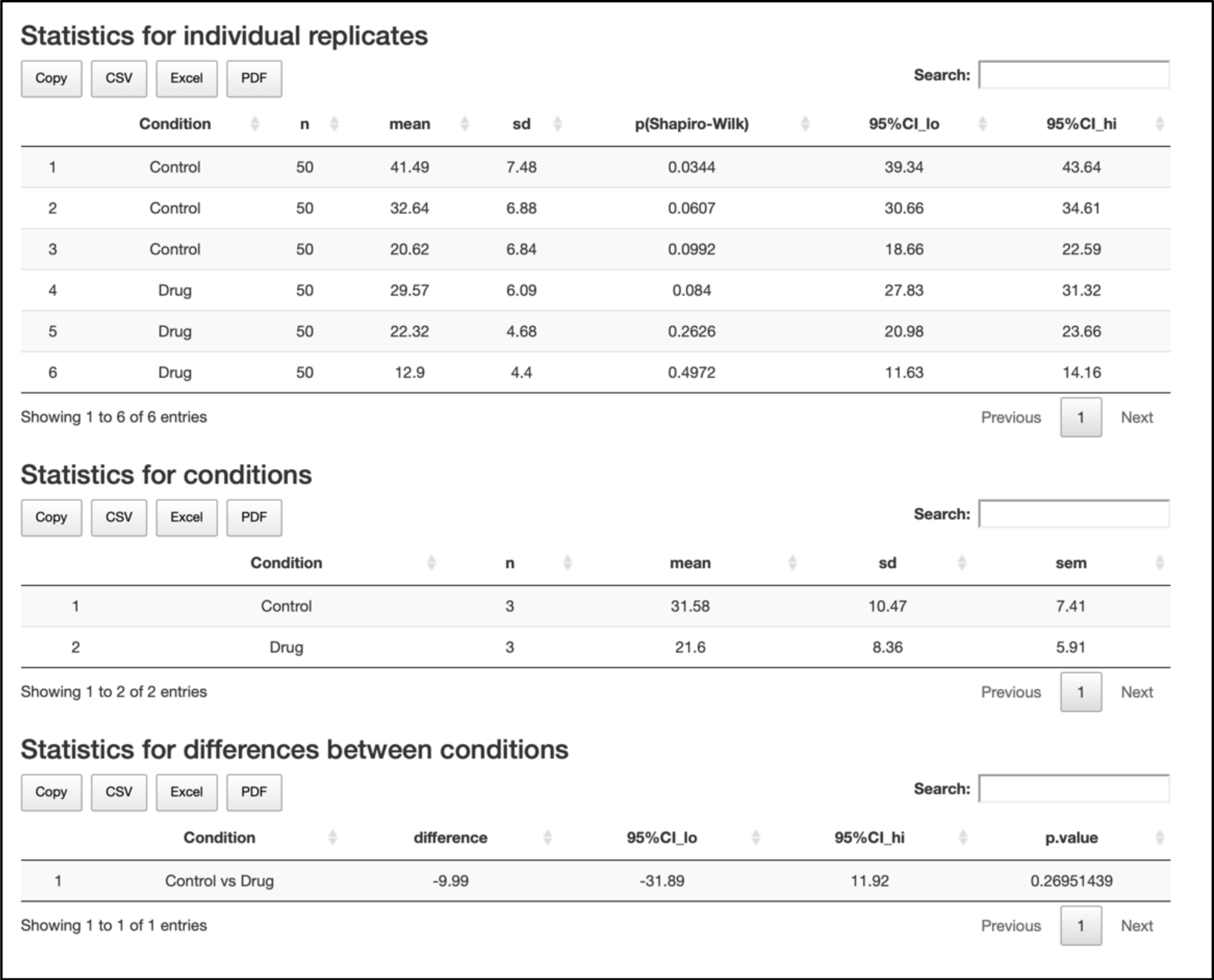
A screenshot of statistics from the example data that are calculated by SuperPlotsOfData. The statistics are available under the ‘Data Summary’ tab. Each of the tables can be downloaded in a number of formats.

### Biological replicates

The statistics for each condition are presented in a second table that is available under the ‘Data Summary’ tab. The number of (biological) replicates, the average, standard deviation and standard error of the mean are displayed.

### Comparing conditions

Depending on the experimental design, the biological replicates can be related. To highlight a paired relation, dashed lines can be added to connect the mean or median values of the replicates. In this situation the data is treated as paired in the statistical analysis.

The aim of an experiment is often to detect a difference between the experimental conditions. The predominant statistical methods for the quantitative comparison of data is a null-hypothesis significance test (NHST). A low p-value provides evidence for a difference between conditions. However, it is often more interesting and biologically relevant to ‘how large is the difference?’ (Cumming, 2014). The difference between conditions and the 95% confidence interval is also known as the effect size and this type of analysis is termed estimation statistics (Claridge-Chang and Assam, 2016). In SuperPlotsOfData there is an option to display the difference between a reference condition that can be selected by the user and the other conditions. A p-value from a student’s t-test is also supplied. A paired t-test is done when the means are connected with lines and an unpaired t-test is done (assuming unequal variances, also known as Welch’s t-test) when the means are not connected. The table with statistics can be displayed under the plot and is also available under the ‘Data Summary’ tab.

## Optimizing the visualization

Scientific graphs often represent data in an unpolished format with default settings that are not optimized to communicate the data in an easy-to-digest manner. In marked contrast, the field of data visualization focusses on ‘storytelling’ and aims to convey the story that is told by the data with a clear and compelling illustration. Several of the principles of storytelling may aid the construction of graphs that are easier to understand. SuperPlotsOfData has several features that improve the plot by making the communication and interpretation of the data more effective, including (i) the option to sort data according to the measured variable, (ii) the choice to rotate the graph by 90 degrees to improve the readability of the conditions, (iii) effective use of colorblind friendly colors, and (iv) the option to switch off the gridlines. Finally, a dark theme is available to generate plots for presentation or websites that use a dark background.

## Output

The graphs can be downloaded in PDF or PNG format. The PDF format enables editing of figures in software applications that accept vector-based graphics. The statistics can be downloaded in CSV or Excel file format.

A snapshot of the current setting can be made by the ‘Clone current setting’ button, which returns a URL with that encodes the user-defined settings of the current session. When the data is imported from a web address, the graph can be stored and exchanged. This option enables a reproducible user interface.

In addition, the option to retrieve a hyperlink of the current setting facilitates data sharing and re-use. For details, the reader is referred to the papers that report our other apps in which this feature is implemented (Postma and Goedhart, 2019; Goedhart and Luijsterburg, 2020). The hyperlink (URL) that corresponds to the setting used in the plots shown in figure 1 and figure 3 are listed in table 1.

**Table 1:** Hyperlinks that define the plots in figures 1 and 3. Launching the webtool by using the hyperlink will reproduce the corresponding figure (make sure to copy-paste the entire URL. Clicking the hyperlink in the pdf may result in a broken link).

| Figure | URL |
| --- | --- |
| 1A | https://huygens.science.uva.nl/SuperPlotsOfData/?data=2 |
| 1B | https://huygens.science.uva.nl/SuperPlotsOfData/?data=2&vis=;;0.7;;;TRUE;1;none&layout=No;;;;;;;;;1 |
| 3A | https://huygens.science.uva.nl/SuperPlotsOfData/?data=2;;Treatment;Speed;Replicate&vis=quasirandom;;0.7;mean;TRUE;;;1;none&layout=No;;;;;;;;;6;;480;480&color=none&label=;;;;;;;;;24;24;18;;& |
| 3B | https://huygens.science.uva.nl/SuperPlotsOfData/?data=2;;Treatment;Speed;Replicate&vis=quasirandom;;0.7;mean;TRUE;;;1;none&layout=Horizontal;;;TRUE;;0,60;;6;;480;480&color=none&label=;;;;;;;;;24;24;18;;& |
| 3C | https://huygens.science.uva.nl/SuperPlotsOfData/?data=2;;Treatment;Speed;Replicate&vis=quasirandom;TRUE;0.7;mean;TRUE;;;1;none&layout=No;TRUE;;TRUE;;0,60;;6;;480;480&color=none&label=;;;;;;;;;24;24;18;;& |

**Figure 3:**
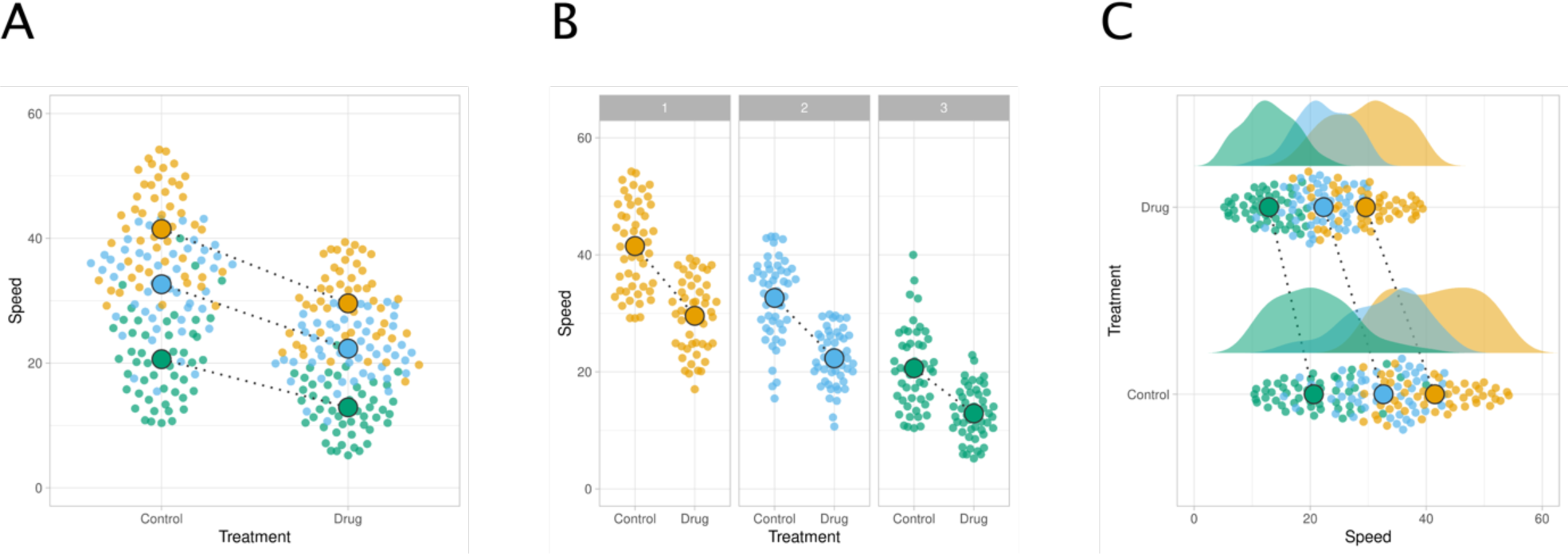
An example of the flexibility in plotting the data that is offered by SuperPlotsOfData. All plots are based on the example data and the biological replicates are paired, as indicated with the dashed line. (A) classic presentation (B) separate presentation of each replicate and (C) rotated plot with de data distributions on top, also known as a rain cloud plot.

## Limitations

### Data format

The web app uses the tidy format as input (Wickham, 2014). This format is substantially different from the spreadsheet, or wide, format that probably most scientists are used to. Although the information content is identical for both formats, it is distributed in a different way over rows and columns. Working with the tidy format and getting the data in the right shape can be challenging for anyone that is not used to the tidy data format.

### Statistical analysis

Experiments often have a ‘nested’ design, consisting of both technical and biological replicates. Each type of replicate contributes in a different way to the overall variance. A statistically correct comparison between conditions, taking the experimental design into account, uses a multilevel model (Aarts et al., 2014; Galbraith et al., 2010). However, this is a relatively complicated analysis. As a practical and intuitive alternative, the average of each technical replicate can be used as input for a standard (paired) t-test. This approach is statistically valid (Galbraith et al., 2010; Aarts et al., 2014), but it is recommended to keep the number of measurements for technical each replicate similar (Lord et al., 2020). Although averaging the technical replicates is not the best approach (Galbraith et al., 2010), it is preferred over aggregating all technical replicates and ignoring the existence of biological replicates. For a detailed discussion about the superplot, the reader is referred to the original paper that proposed superplots (Lord et al., 2020).

## Conclusion

The SuperPlotsOfData app implements some of the recent innovations in data visualization and analysis. We hope that the webtool will encourage users to adopt best practices in data presentation and analysis. These best practices include the display of individual observations, distinguishing between technical and biological replicates and the use of estimation statistics for the quantitative comparison of conditions.

The tool democratizes modern data visualization as (i) the app is freely accessible online or it can run locally with free software, (ii) it does not require any coding skills and (iii) it has a minimal learning curve. Finally, the code of the app is available which makes the analysis procedure transparent and open to modification to accommodate any future developments in analysis and visualization.

## Acknowledgments

A good chunk of the code for SuperPlotsOfData is taken from a number of previously developed apps (https://github.com/JoachimGoedhart); PlotsOfData, VolcaNoseR, and PlotsOfDifferences, which in turn benefit from code that is shared on repositories (Github), blogs and fora (stackoverflow). We thank Auke Folkerts (UvA, The Netherlands) for help with the server that runs shiny. The enthusiasm and support from colleagues on Twitter are highly appreciated and the interaction with Samuel J. Lord was particularly helpful for the implementation of the superplot.

## Data availability

All data and code is available in public repositories (GitHub and Zenodo) as referenced in the manuscript.

